# Scaling of Internal Organs during *Drosophila* Embryonic Development

**DOI:** 10.1101/2020.12.15.422810

**Authors:** P. Tiwari, H. Rengarajan, T. E. Saunders

## Abstract

Many species show a diverse range of sizes; for example, domestic dogs have large variation in body mass. Yet, the internal structure of the organism remains similar, *i.e*. the system scales to organism size. *Drosophila melanogaster* has been a powerful model system for exploring scaling mechanisms. In the early embryo, gene expression boundaries scale very precisely to embryo length. Later in development, the adult wings grow with remarkable symmetry and scale well with animal size. Yet, our knowledge of whether internal organs initially scale to embryo size remains largely unknown. Here, we utilise artificially small *Drosophila* embryos to explore how three critical internal organs – the heart, hindgut and ventral nerve cord (VNC) – adapt to changes in embryo morphology. We find that the heart scales precisely with embryo length. Intriguingly, reduction in cardiac cell length, rather than number, appears to be important in controlling heart length. The hindgut – which is the first chiral organ to form – displays scaling with embryo size under large-scale changes in the artificially smaller embryos but shows few hallmarks of scaling within wild-type size variation. Finally, the VNC only displays weak scaling behaviour; even large changes in embryo geometry result in only small shifts in VNC length. This suggests that the VNC may have an intrinsic minimal length, which is largely independent of embryo length. Overall, our work shows that internal organs can adapt to embryo size changes in *Drosophila*. but the extent to which they scale varies significantly between organs.

## Introduction

Organism scaling (allometry) – including in humans – is an important developmental process [1]. A striking example is the large variation due to selective breeding in domestic dog size, from ~2-3kg to ~70kg, yet they are genetically very similar [2]. The extent to which an organ grows is regulated by the body size of the individual. Understanding the processes that drive organism scaling has been a longstanding challenge in developmental biology [3, 4]. In recent years, the study of external organs, such as the insect wing [5–7] and beetle horns [8], has provided novel insights into the relationship between organ and organism size, and the associated physiological properties [9].

What are the mechanisms controlling organ size? In the case of winged organisms, they require extremely precise regulation of the wing size, including left/right symmetry, to ensure optimal flying [10]. Such regulation could be driven by both genetic and mechanical processes within the organ [11, 12]. In *Manduca sexta*, wing size is tightly regulated by the adult body size. This regulation is achieved through altering the cell number in wing disc [5]. Mechanical processes can act through direct limitation on tissue growth or by inducing mechanosensitive pathways [13–17]. The timing of signal interpretation and external hormonal inputs can also direct organ scaling [6, 18–20] and neighbouring organs can regulate organogenesis [21]. Theoretical approaches have also been important in deciphering the possible processes driving precise scaling [22].

Arguably the best quantified system for studying scaling in development is gene expression boundary specification along the anterior-posterior (AP) axis in the early *Drosophila* embryo [23]. Such boundaries scale precisely with embryo length (E_L_), with an error of less than one cell diameter. These boundaries are downstream of the morphogen gradient Bicoid and other maternal inputs [24, 25]. The scaling of the boundaries is robust to natural variations in both Bicoid levels and embryo size, but the scaling does breakdown under larger changes in either Bicoid or embryo size [26, 27].

Later in *Drosophila* development, there has been extensive research into how larval organs grow and adapt to environmental stresses [28–30]. These organs form first in the embryo and are derived from the precisely scaled patterns laid down in the early embryo [31]. During larval and pupal stages, most of these organs undergo substantial reconfiguration [30] including large-scale removal of the initial embryonic cells. So, whether these initial embryonic organs need to scale and adjust to changes in embryo size is an open question. Dissecting organ scaling is complicated by the diverse range of organ formation processes and morphologies. For example, in *Drosophila*, muscle growth occurs through myoblast fusion and elongation [32], the hindgut grows by chiral reorientation of cells [33], the ventral nerve cord (VNC) undergoes large scale elongation followed by condensation [34, 35] and the heart forms from a fixed number of cells [31, 36]. Here, taking advantage of imaging accessibility of the *Drosophila* embryo and a mutant that produces smaller (yet still viable) embryos, we provide quantitative measures of the formation and scaling of three internal organs: the heart, hindgut, and VNC. Intriguingly, they all display distinct scaling characteristics, suggesting that embryonic internal organs are highly variable in how they respond to changes in organism size.

## Methods

### Generation of artificially smaller embryos

In wild-type (control) embryos, the natural variation is approximately 10% in embryo length (Fig. 1A, B) and 13% in embryo width (Fig. 1A, C). This small variability makes studying scaling within the embryo challenging. To get around this problem, we utilised *TrafficJam-Gal4>UAS-fat2-RNAi* (henceforth referred to as *TjGal4>fat2RNAi)* expressing flies [27, 37], which lay shorter embryos along the anterior-posterior (AP) axis, though they have larger width (Fig. 1A-C). Embryos labelled as *TjGal4>fat2RNAi* are laid by females of the genotype *TjGal4>fat2RNAi*. Importantly, these eggs are viable and typically result in healthy larvae (Movie S1). This manipulation provided 30% variation in the embryo length and 20% variation in the embryo width (E_W_). The control embryos have a length distribution of 473 μm to 572 μm and width distribution of 150 μm to 195 μm. In the *TjGal4>fat2RNAi* embryos we observe length distribution of 324 μm to 465 μm and width distribution of 181 μm to 228 μm. Embryo length is correlated with embryo width in the *TjGal4>fat2RNAi* embryos, but not in control (Fig. 1D). Comparing the embryo length with the aspect ratio (length/width) we see a clear correlation in controls and *TjGal4>fat2RNAi* embryos (Fig. 1E). As reported previously [27], this corresponds to the *TjGal4>fat2RNAi* embryos only have a small decrease in total volume compared with wild-type embryos.

**Figure 1.**
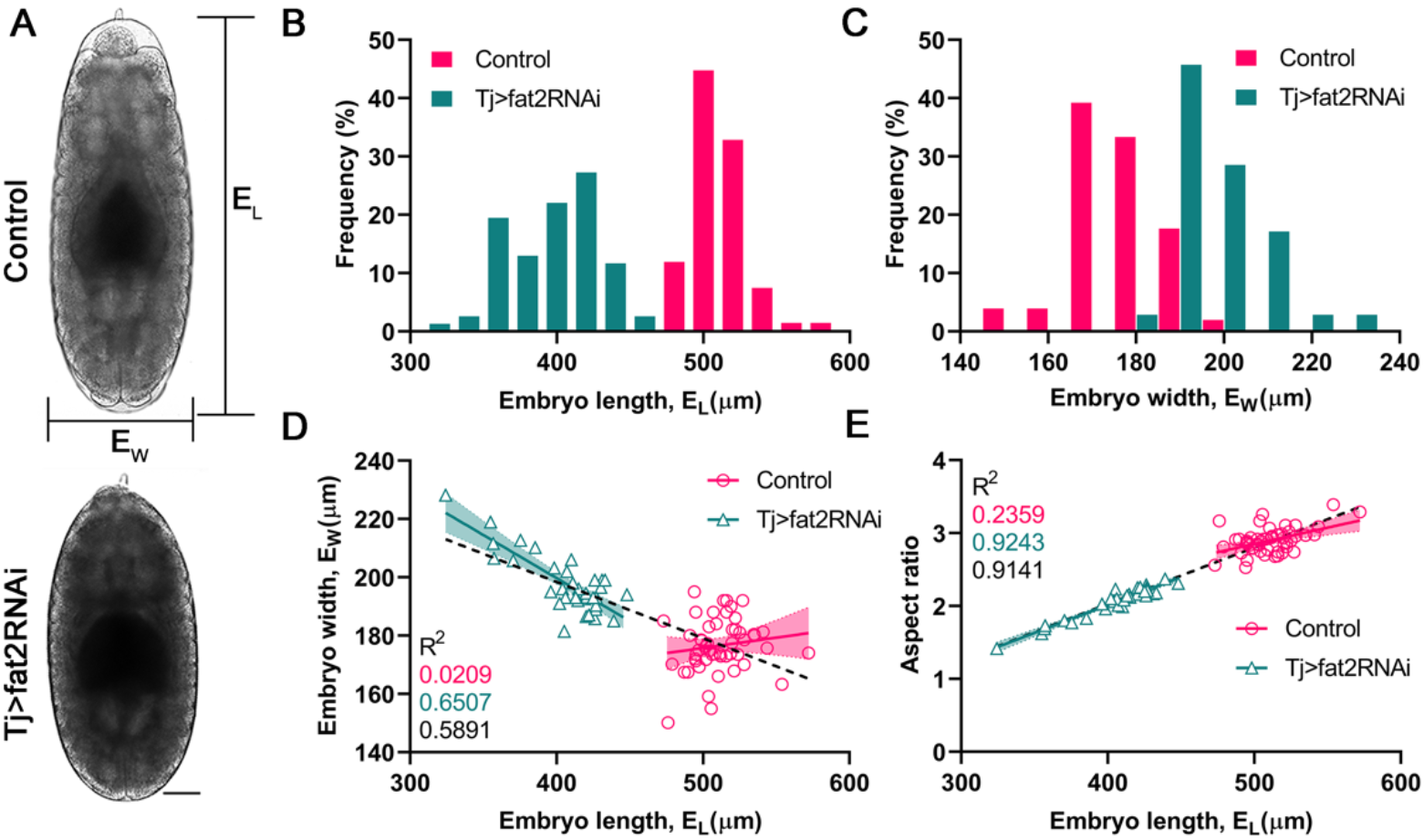
Quantification of embryo morphology in control and TjGal4>fat2RNAi embryos. A) Brightfield images of stage 14 wild-type (control) and *TjGal4>fat2RNAi* embryos on same scale. B) Distribution of embryo length in control and *TjGal4>fat2RNAi* embryos. C) Distribution of embryo width in control and *TjGal4>fat2RNAi* embryos. D) Correlation of embryo length with embryo width in control and *TjGal4>fat2RNAi* embryos. E) Correlation of aspect ratio with embryo length. For (B), n=67 (control), n=77 (*TjGal4>fat2RNAi*), for (C-E) n=51 (control), n=35 (*TjGal4>fat2RNAi*). Scale bar in (A) is 50 μm. Shaded regions represent 95% confidence interval on the linear fitting and r^2^ values given in the legend, colour coded by control (magenta), *TjGal4>fat2RNAi* (cyan) and all data (black).

### Fly stocks

We used the following fly lines in this paper: *Tj*Gal4 [38], UAS-*fat2*RNAi (VDRC27114 and BDSC40888), Hand::GFP and *Hand*-Gal4 [39], UAS-moe::GFP [40], *elav*Gal4>UAS GFP (BDSC5146), *Byn*Gal4>UAS-myr::GFP [41], *Byn*::mtdTomato (from Kenji Matsuno), *Histone::RFP* (BDSC 23650), *Myo31DF^L152^; Byn*Gal4>UAS-myr::GFP[41]. In the heart experiments, our control line was Hand::GFP. In the VNC experiments, our control line was *elav*Gal4>UAS GFP. In the hindgut experiments, our control line was *Byn*::mtdTomato. We confirmed that *Byn*>Gal4-UAS>myr::GFP UAS-*fat2*-RNAi showed similar behaviour to our control embryos for the hindgut.

### Imaging protocols / microscope

We used NikonA1R and Zeiss LSM 700 confocal microscopes for time lapse imaging. We picked the desired staged embryos from the apple agar plate and dechorionated using bleach. Embryos were then washed in PBS and mounted on the coverslip dish in the desired orientation (dorsally for heart and hindgut and laterally for VNC). Time lapse imaging was performed on a confocal microscope with time interval of 5-10 minutes. Recordings were taken at room temperature, typically 21°C to 23°C. All data is collected from live movies except Movie S5, which is immunostained for GFP and DE-Cad following standard immunostaining protocols. Chick anti-GFP (1:10000 abcam) and Rat-DCAD2 (1:300, DSHB) primary antibodies were used to label the GFP expressing cells and DE-Cad expression respectively. The primary antibodies were detected with Alexaflour labelled secondary antibodies (1:400, Invitrogen).

### Image analysis details

Embryo length and width were measured in ImageJ manually as shown in Fig. 1A. *Heart quantification*. At stage 14, we measured the two rows of cardioblasts once they clearly move inwards from a dorsal view. For each embryo we measured both rows of cardioblasts and averaged the values for each embryo (Fig. 2A-C). For each embryo we measured the distance between the anterior-most and posterior-most ends of both rows and averaged these values. We used this value to measure the heart arching at stage 14. At stage 16, we measured the size of the heart immediately after both rows of cardioblasts met at the dorsal midline. *VNC quantification*: VNC length measurements were done by drawing a segmented line along the VNC in ImageJ at two developmental time points (Fig. 4A-B): the onset of stage15 (we used the gut morphology to identify this specific landmark); and during stage 17, just prior to clear tracheal filling. *Hindgut quantification*: Hindgut measurements (Fig. 5A-B) were done at the developmental time point when the hindgut reached its maximum distance along AP axis before twisting (during stage 14). Distances along the AP axis were measured from the anus in the posterior manually on ImageJ. Curvature was measured by fitting a circle to the region highlighted in Fig. 5B. As the hindgut anterior end is not perfectly circular in its morphology, we fitted the circle to the highlighted dark blue region in Fig. 5B. For each data set, all embryos within the cohort were quantified prior to any statistical analysis to reduce potential for bias. Only embryos that displayed clear evidence of sickness were excluded from the analysis.

**Figure 2.**
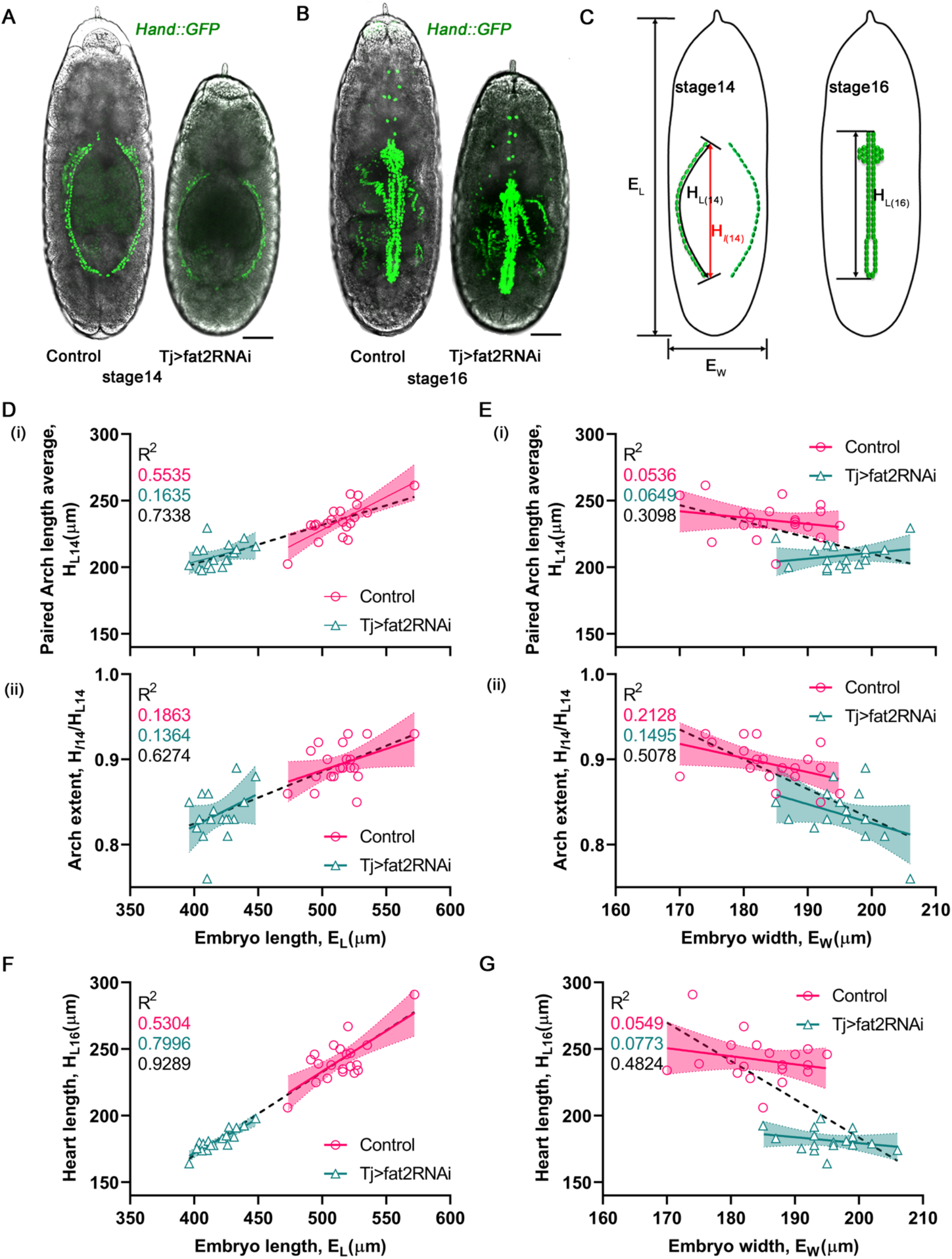
Scaling of the Drosophila embryonic heart. A) Two contralateral opposite rows of heart cells at stage 14 in control and *TjGal4>fat2RNAi* embryos marked by Hand::GFP (Methods). B) Stage 16 heart cells align close to each other to form the heart tube [45]. C) Schematic showing different measurements used in analysis: H_*L(14)*_ and H_*L(16)*_ are the total length of heart at stage 14 and 16 respectively; and H_*l14*_ is the linear distance between the anterior and posterior end of the row of cells at stage 14. D) Correlation with embryo length at stage 14 of average total heart length, H_*L(14)*_ (i) and arch extent, H_*l(14)*_ /H_*L(14)*_ (ii). E) Correlation with embryo width at stage 14 of average heart length at stage 14 (i), and arch extent (ii). F-G) Correlation of heart length at stage 16, H_*L(16)*_, with embryo length (F) and width (G). For control embryos (n=20) and for *TjGal4>fat2RNAi* embryos (n=17). Shaded regions represent 95% confidence interval on the linear fitting and r^2^ values given in the legend, colour coded by control (magenta), *TjGal4>fat2RNAi* (cyan) and all data (black).

### Statistical analysis

All statistical analysis was performed in GraphPad except for the Bootstrapping analysis (Fig. 4E). For calculating the p-value between measured means we used Welch’s test (unpaired t test with Welch’s correction). To test whether observed scaling relationships showed statistically significant differences from no correlation (null hypothesis that zero gradient in the linear fit) we used GraphPad’s built-in regression-slope test. We further provide r^2^ values on all linear fittings in the figure panels. In all linear regression analyses, we show the 95% confidence interval (shaded regions in figure panels). In Fig. 4E, error on the VNC embryo-to-embryo variability was calculated through Bootstrapping using the Matlab function *bootstrp*. 100 random samples were generated for each condition.

## Results

### The *Drosophila* heart scales with embryo length

The *Drosophila* heart is a tubular structure with no change in cell number during its formation [42]. The two rows of cells (52 cells each) migrate from the two lateral sides of embryo (at developmental stage 14) towards the dorsal midline (Movie S2). They match precisely at the dorsal midline in developmental stage 16 to form the dorsal vessel [43, 44]. In stage 14, the two rows of cells arrange in arched structures (Fig. 2A). In stage 16, the heart has a simple linear structure (Fig. 2B). From imaging embryos expressing heart-specific markers (Methods) we can quantify the heart shape during morphogenesis in different sized embryos (Fig. 2C).

At stage 14, we see that the future heart in *TjGal4>fat2RNAi* embryos is shorter in total length (H_L14_ = 209±9 μm) than control embryos (H_L14_ = 235±14 μm) (Fig. 2D-E). Further, the heart is more arched in *TjGal4>fat2RNAi* embryos compared to stage 14 control embryos (Fig. 2D(ii)). At stage 14 in control embryos, the length of the future heart sections scales with the embryo length (p < 10^-3^) but not with the embryo width (p = 0.33) (Fig. 2D-E). In the *TjGal4>fat2RNAi* embryos at stage 14 there is not significant scaling of heart length with the embryo size (p = 0.11 and 0.32 for embryo length and width respectively). We define the heart arching as H_*l14*_/H_*L14*_; a straight heart would have an arch of 1, with the value of H_*l14*_/H_*L14*_ decreasing for more curved hearts. There is a weak scaling between the heart arch extent in stage 14 and embryo length within each data set (p = 0.04 for control embryos, Fig. 2E(ii)) but when combining the data sets, we see a clear trend for increasingly arched hearts at stage 14 in shorter, wider embryos (Fig. 2E(ii)).

At stage 16, control and *TjGal4>fat2RNAi* embryos show clear scaling with embryo length (p<10^-3^ for both conditions, Fig. 2F). The correlation between embryo and heart length is substantially increased compared with stage 14 embryos. This suggests that the heart length adjusts to embryo size during its formation. This strong scaling is remarkable for two reasons: (1) the heart is constructed of a fixed cell number; and (2) the scaling becomes more apparent later in embryonic development. Given the linear nature of the heart orientated along the AP-axis, unsurprisingly, we do not see scaling of heart size with embryo width (Fig. 2G).

### The embryonic *Drosophila* heart scales by reducing cell length along the AP axis

Given the clear scaling of the stage 16 heart with embryo length, we asked what morphological changes occur to ensure such scaling? Two possibilities are that the cardioblasts change morphology or that the number of heart cells is reduced. To count the number of cardioblasts, we used Hand::GFP, which marks the nuclei of the cardioblasts and surrounding pericardial cells, which are readily distinguished by position (Fig. 3A). We counted the number of heart cells in control and *TjGal4>fat2RNAi* embryos.

**Figure 3.**
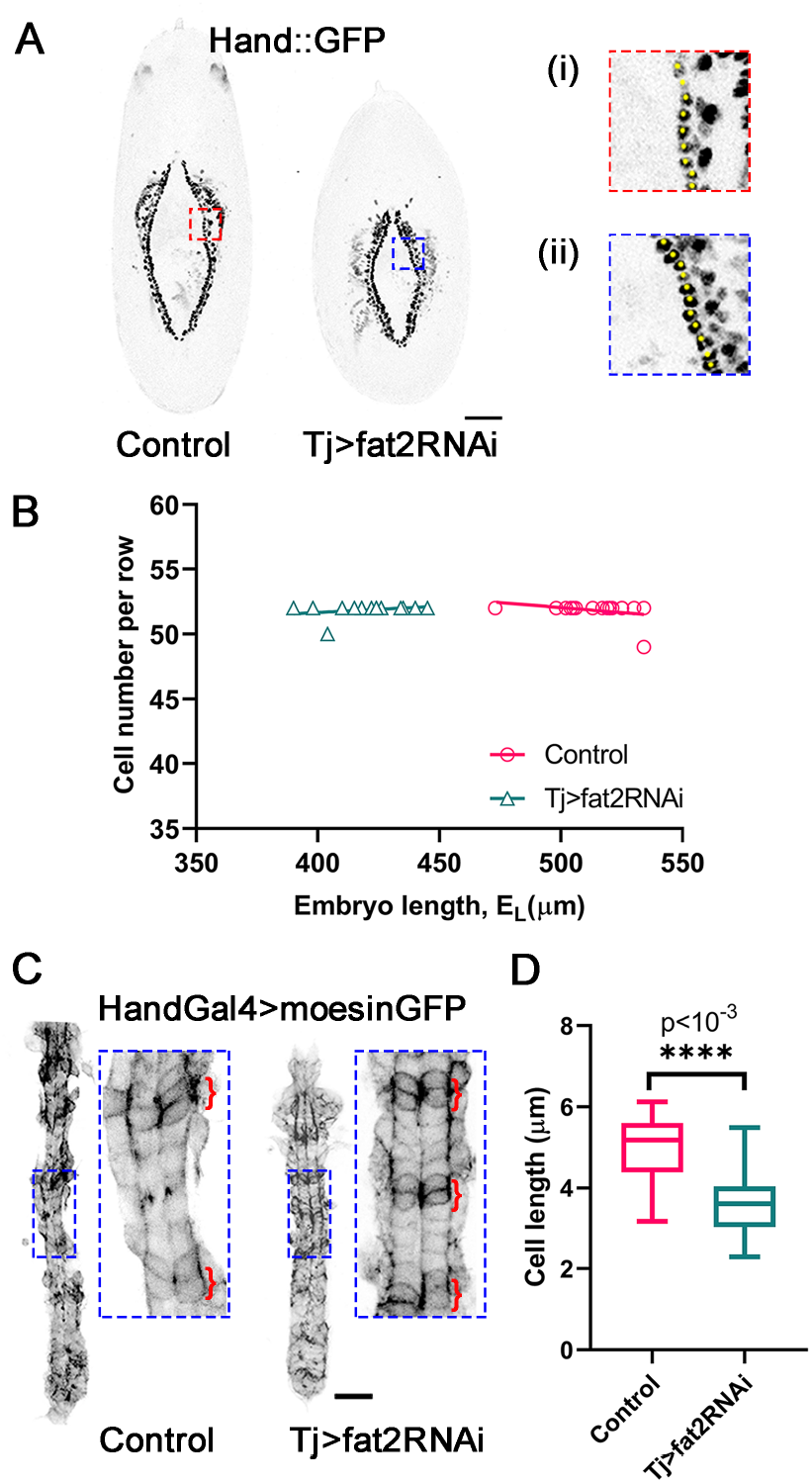
Scaling of the embryonic heart occurs through cell shape changes. A) Heart cells (cardioblasts) with their nuclei labelled with Hand::GFP. (i) and (ii) corresponds to zoomed in regions denoted by the red and blue boxes on left respectively; cardioblasts marked with yellow dots. (B) Cardioblast number against embryo length for control and *TjGal4>fat2RNAi* embryos, n=17 (control), n=15 (*TjGal4>fat2RNAi*). C) *Hand*-Gal4>moe::GFP marking the cell boundary in the developing heart for control (left) and *TjGal4>fat2RNAi* (right) embryos. Red brackets highlight Seven-Up positive cells which are narrower in the AP-axis than Tinman-positive cardioblasts. The boxed regions are the same absolute size. D) Mean cardioblast length in AP-axis in control and *TjGal4>fat2RNAi* embryos (**** p < 10^-3^). Box represents 25%-75% percentiles and error bars are standard deviation. Lines in (B) represent a linear fitting with r^2^ values given in the legend, colour coded by control (magenta) and *TjGal4>fat2RNAi* (cyan).

We found that the number of cardioblasts on each side of the embryo remained constant, with 52 per embryo, even in very short embryos (Fig. 3B), showing that heart scaling is independent of cell number. To test whether the cell size along the AP axis was altered, we imaged cardioblast morphology using *Hand*-Gal4>UAS-Moe::GFP, which marked the cell boundaries (Fig. 3C). We observed a clear decrease in the cell length along the AP-axis, with 5.0±0.7μm (control) and 3.6±0.7 μm (*TjGal4>fat2RNAi*), p<10^-3^ (Fig. 3C-D). The heart is comprised of a repeating pattern of four Tinman-positive and two Seven-up (Svp)-positive cardioblasts [45]. The Svp-positive cardioblasts form ostia and are generally narrower than the Tinman-positive cells. In Fig. 3C, we highlight the Svp-positive cardioblasts (red brackets). Within the same spatial region, there are two sets of Svp-positive cardioblasts in the control embryo compared to three sets in the *TjGal4>fat2RNAi* embryos. Overall, we see that by stage 16 the heart scales with embryo length and this is mediated by a change in heart cell morphology, not number.

### The Ventral Nerve Cord weakly adjusts to embryo size changes

The VNC, a part of the *Drosophila* Central Nervous System, is first specified ventrally during germband elongation (stage 8 onwards). By stage 11, the developing VNC along with the germ band extends to the dorsal side. Then, from stage 12 onwards it retracts along with the germband. By stage 17, it has reduced in size by 60% [31, 35] and resides near the ventral surface (Fig. 4A and Movie S3). During the condensation process, a significant number of cells die through apoptosis [46]. We measured VNC length at stage 15 and late stage 17 (Methods) in control and *TjGal4>fat2RNAi* embryos and compared it to the embryo length. At stage 15 the total VNC length V_L15_ (Fig. 4B) was 442±13 μm and 403±16 μm, for control and *TjGal4>fat2RNAi* embryos respectively (for control (n=16) and *TjGal4>fat2RNAi* (n=34), p<10^-3^, difference in means). The total VNC length in *TjGal4>fat2RNAi* embryos was around 10% shorter than in control (p <10^-3^, difference in means). In stage 17, the total VNC length in *TjGal4>fat2RNAi* embryos was also around 10% shorter than control conditions (267±23 μm and 235±14 μm for control (n=16) and *TjGal4>fat2RNAi* (n=23), p<10^-3^, difference in means).

**Figure 4.**
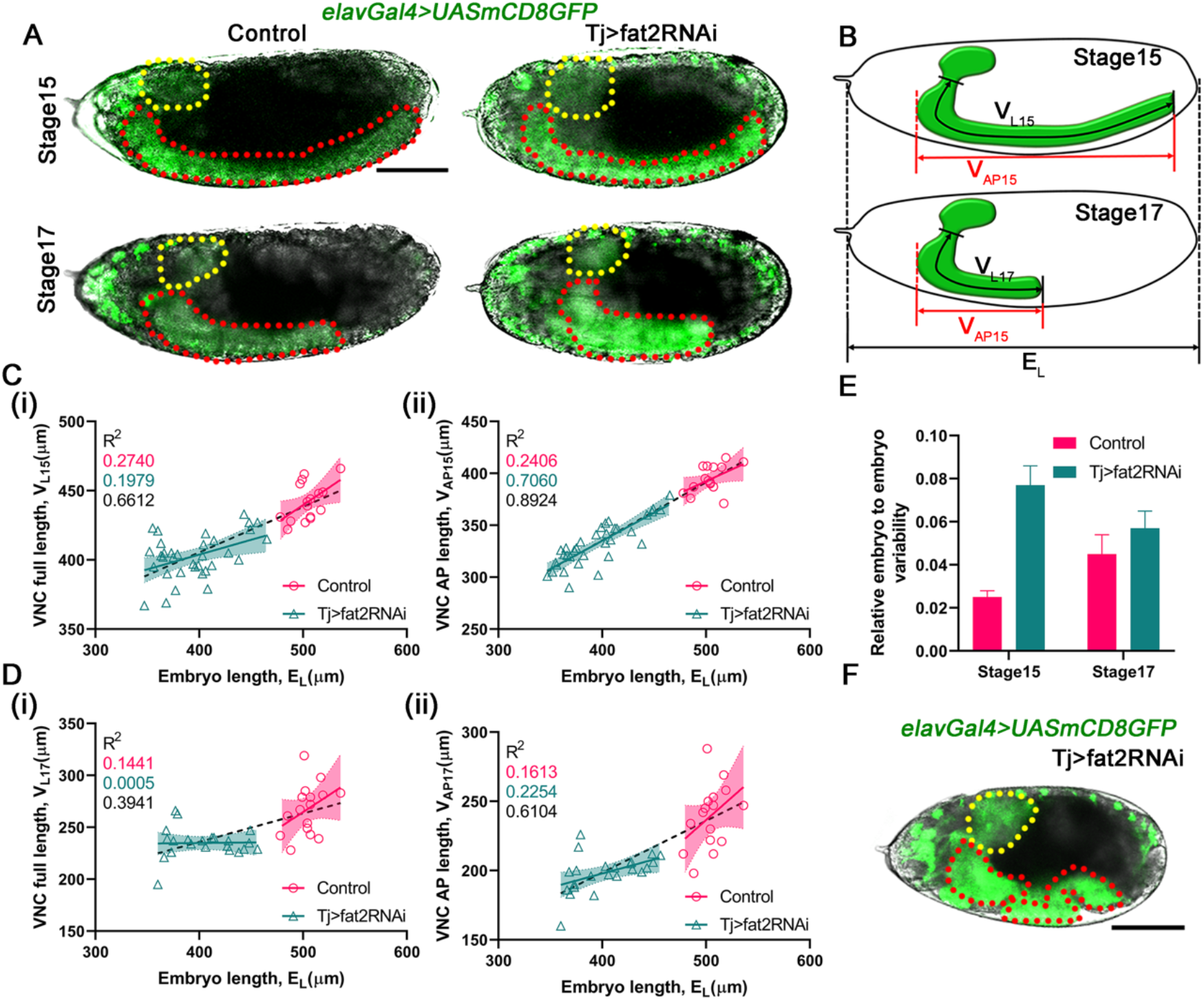
Scaling of the embryonic VNC. A) Stage 15 and 17 control and *TjGal4>fat2RNAi* embryos (CNS labelled in green by *elav*Gal4>UAS-GFP). Red and yellow dotted regions mark the VNC and brain respectively, B) Schematic showing the measurement method for the VNC length analysis, V_L_ is the length from the neck region of CNS to the posterior end of CNS, V_AP_ is the length of VNC along the AP axis, E_L_ is the embryo length on the AP axis, numbers (15/17) in the subscript denote stage. C-D) Correlation between total VNC length (i) and VNC length along AP-axis (ii) with embryo length at stage 15 (C) and stage 17 (D). E) Embryo-embryo variation in VNC length / embryo length at stages 15 (n = 16, 34 respectively) and 17 (n = 16, 23 respectively) for control and *TjGal4>fat2RNAi* embryos. Error bars found by Bootstrapping (Methods). F) Example of buckling occurring in the VNC in a *TjGal4>fat2RNAi* embryo. All scale bars = 100 μm. Shaded regions represent 95% confidence interval on the linear fitting and r^2^ values given in the legend, colour coded by control (magenta), *TjGal4>fat2RNAi* (cyan) and all data (black).

The total VNC length correlated with embryo length at stage 15 in both control and *TjGal4>fat2RNAi* embryos (Fig. 4C(i)). Interestingly, the linear length of the VNC along the AP-axis (V_AP15_) scaled more strongly with embryo length in the *TjGal4>fat2RNAi* embryos (Fig. 4C(ii)). However, by stage 17 this correlation was largely lost in both control and *TjGal4>fat2RNAi* embryos (Fig. 4D). In particular, despite the large changes in embryo geometry (embryo length between 347 μm and 465 μm), in *TjGal4>fat2RNAi* embryos we saw no clear correlation between the VNC total length and the embryo length (Fig. 4D(i)). As with stage 15, there is slightly stronger scaling with embryo length when we analyse the linear length of the VNC along the AP-axis (Fig. 4D(ii)). When we compare the mean total VNC length in control and *TjGal4>fat2RNAi* embryos in stage 17 we do see a small change in length, but this is less than 10% of the mean VNC length (compared to ~30% shortening in embryo length). These results suggest that during the final stages of VNC condensation (stages 16 and 17), the VNC length only weakly adapts to the surrounding embryonic environment.

This behaviour motivated us to explore the variability in VNC length during development. Looking at the variation in VNC length in control conditions (n= 16 embryos), we saw that the VNC length is highly robust between embryos, even though the embryos have variable length (478 μm to 536 μm) (Fig. 4E). The relative error in the VNC length was less than 4% in control embryos. The variability was larger in *TjGal4>fat2RNAi* embryos, but still less than 10% variation. We also found the absolute error in the total VNC length (normalised by average VNC length) – the VNC length variation between embryos was 3% and 4% for control and *TjGal4>fat2RNAi* embryos respectively in stage 15, and 9% and 6% for control and *TjGal4>fat2RNAi* embryos respectively in stage 17. The embryo-to-embryo variability in VNC length was less than 10% of its typical length in both control and *TjGal4>fat2RNAi* embryos, regardless of embryo length. This is remarkably small variation for an organ that undergoes such large-scale morphological changes. These results suggest that the final total VNC length may have a preferred intrinsic length and perhaps even a minimum size requirement (>220μm in stage 17), though further work is required to test these ideas. A prediction stemming from these observations is that in very short embryos the VNC may deform, as there could be insufficient space for it to occupy the ventral surface. In one *TjGal4>fat2RNAi* embryo (which displayed muscle twitching but did not hatch), we indeed observed buckling (Fig. 4F). However, this buckling was not just in the VNC, so it is not possible to conclude clearly whether the VNC induced buckling or if it was deforming due to other size pressures.

### Scaling in the *Drosophila* hindgut, an asymmetric organ

The *Drosophila* hindgut is an asymmetric organ that grows largely by cell shape change [47] and not proliferation after stage 10 [48]. To explore scaling of the *Drosophila* hindgut, we chose to examine the shape of the gut in stage 14. At this point, the hindgut forms a characteristic inverse “question mark” shape (Fig. 5A and Movie S4). Given the chiral nature of the hindgut, we measured the organ curvature as well as its length and width in both control and *TjGal4>fat2RNAi* embryos (Fig. 5B).

**Figure 5.**
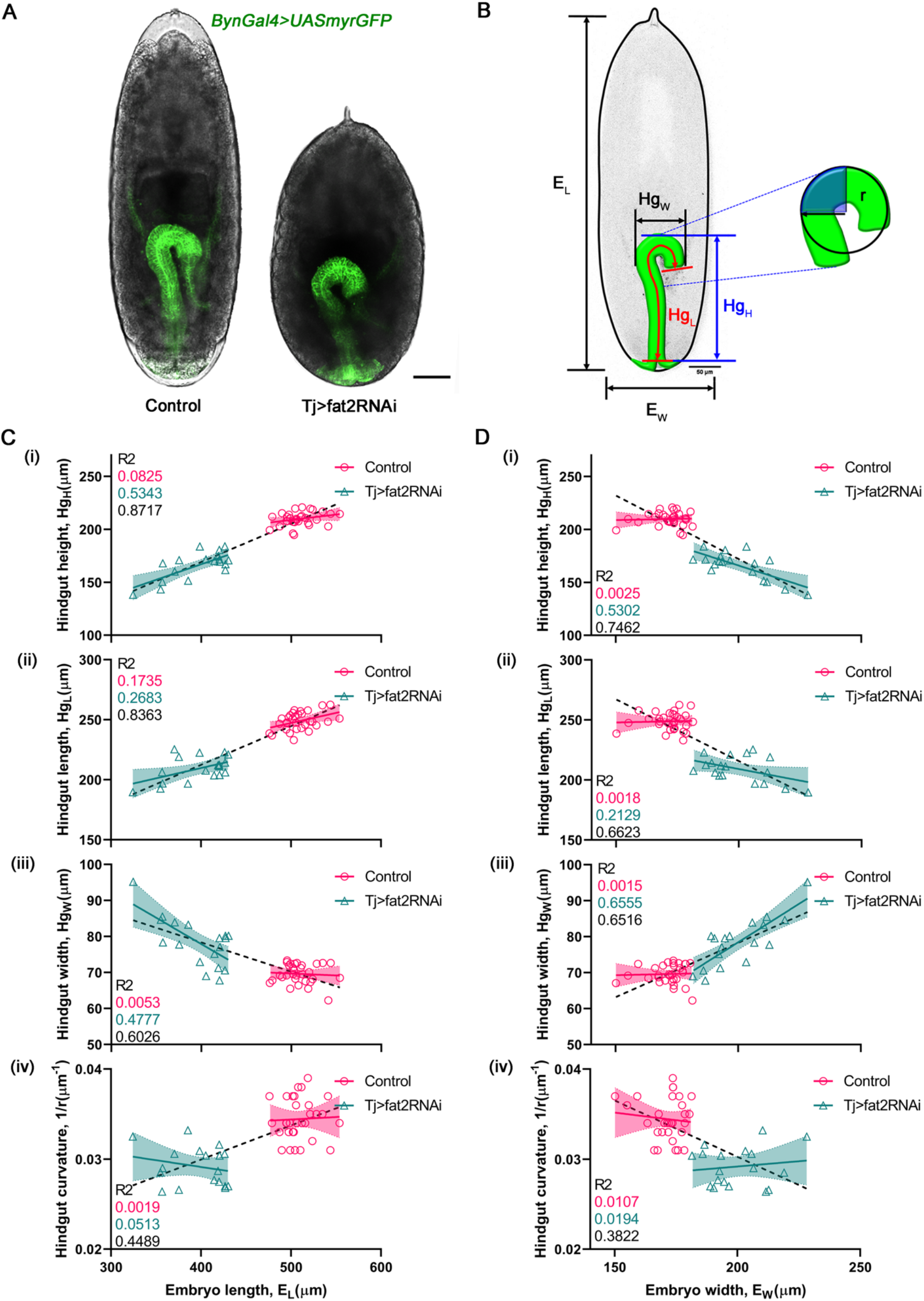
Hindgut scaling at stage 14. A) Representative images of control (left) and *TjGal4>fat2RNAi* (right) embryos expressing *Byn*-Gal4>UAS-myr::GFP. Scale bar = 50μm. B) Schematic of measures used to quantify hindgut morphology. Inset shows how curvature was measured; we used the blue-shaded region to find the curvature as the anterior end of the hindgut was not exactly circular. C-D) Scaling of hindgut height Hg_H_ (i), total length Hg_L_ (ii), maximum width Hg_W_ (iii) and curvature (1/r) (iv) with embryo length (C) and width (D) in control (magenta) and *TjGal4>fat2RNAi* (cyan) conditions. Shaded regions represent 95% confidence interval on the linear fitting and r^2^ values given in the legend, colour coded by control (magenta), *TjGal4>fat2RNAi* (cyan) and all data (black) (n=31 for control and n=18 for *TjGal4>fat2RNAi*).

Measuring the total hindgut length (Hg_L_) and its linear extent along the AP axis (Hg_H_), we saw a clear decrease in the mean value for both Hg_L_ and Hg_H_ between wild-type (Hg_L_ 249±7 μm and Hg_H_ 210±6 μm) and *TjGal4>fat2RNAi* (Hg_L_ 209±11 μm and Hg_H_ 166±13 μm) embryos (p<10^-3^ for difference in means for both Hg_L_ and Hg_H_). Looking at the correlation of Hg_L_ and Hg_H_ with embryo length (Fig. 5C(i-ii)), we saw only very weak scaling in the control embryos (p = 0.02 for Hg_L_ and 0.12 for Hg_H_), but stronger scaling in the *TjGal4>fat2RNAi* embryos (p = 0.03 for Hg_L_ and 0.001 for Hg_H_). We also looked at how the hindgut width and curvature altered with embryo length. We observed distinct differences in hindgut width and curvature between control (width = 70 ± 3 μm and curvature = 0.034±0.002 μm^-1^) and *TjGal4>fat2RNAi* (width = 78 ± 7 μm and curvature = 0.029±0.002 μm^-1^) embryos. The hindgut width inversely scaled with embryo length in the *TjGal4>fat2RNAi* embryos (Fig. 5C(iii), p=0.70 and 0.002 for control and *TjGal4>fat2RNAi* embryos respectively). This is likely related to the *TjGal4>fat2RNAi* embryos displaying an inverse relationship between embryo length and width (Fig. 1). Though the curvature decreased in *TjGal4>fat2RNAi* embryos, it did not show a clear scaling with embryo length (Fig. 5C(iv)).

We saw similar behaviour when looking at the geometric properties of the hindgut in comparison with embryo width (Fig. 5D). In control embryos, there was negligible adjustment of hindgut morphology to changes in embryo width. However, in the *TjGal4>fat2RNAi* embryos, we saw more clear evidence for scaling (p<10^-3^) of Hg_H_ and Hg_W_. This is possibly a consequence of the larger scale morphological differences in the size of the eggshell in these embryos; both length (reduced) and width (increased) change considerably (Fig. 1), though total egg volume only decreased slightly. Overall, these results suggest that, within wild-type variation in embryo size, the hindgut does not adjust significantly to embryo size variation. However, in the more stressed conditions provided by the *TjGal4>fat2RNAi* embryos the hindgut can adjust to the changing physical environment. In particular, the increased width of *TjGal4>fat2RNAi* embryos appears to alter the shape of the hindgut; the width of the hindgut scales with embryo width in *TjGal4>fat2RNAi* embryos, but the curvature does not adjust linearly with embryo length or width.

To examine how the mechanical properties of the hindgut affect its ability to shape and whether this affects scaling, we analysed *Myo1D^-/-^* embryos. Myo1D is a non-canonical Myosin that plays a role in chirality formation, with such embryos often displaying an inverted hindgut [33] (Fig. 6A). Due to genetic limitations, we could not explore the effects of loss of Myo1D in *TjGal4>fat2RNAi* embryos. Instead, we focused on wild-type sized embryos, which had around 20% variation in embryo size. The overall dimensions of the hindgut remain similar in *Myo1D^-/-^* embryos compared to control embryos despite the hindgut inversion. However, there was a much larger variation in hindgut morphology in *Myo1D^-/-^* embryos (as shown by spread of 95% confidence intervals shown in Fig. 6B-C). Note, we only analysed *Myo1D^-/-^* embryos that showed a clear hindgut inversion. Further, the curvature of the hindgut appears more sensitive to changes in embryo morphology (Fig. 6B(iv)). *Myo1D^-/-^* embryos show slightly decreased curvature compared with control embryos (control = 0.034±0.002, Myo1D = 0.032±0.003, p <10^-2^, Fig. 6C(iv)). As with our control embryos, the *Myo1D^-/-^* embryos showed no clear hallmarks of scaling with embryo length or width. We conclude that disruption of Myo1D results in more than simply inverting the hindgut shape; there is much larger embryo-to-embryo variability between embryos and the morphology of the hindgut is not simply an inversion of the “question-mark” geometry.

**Figure 6.**
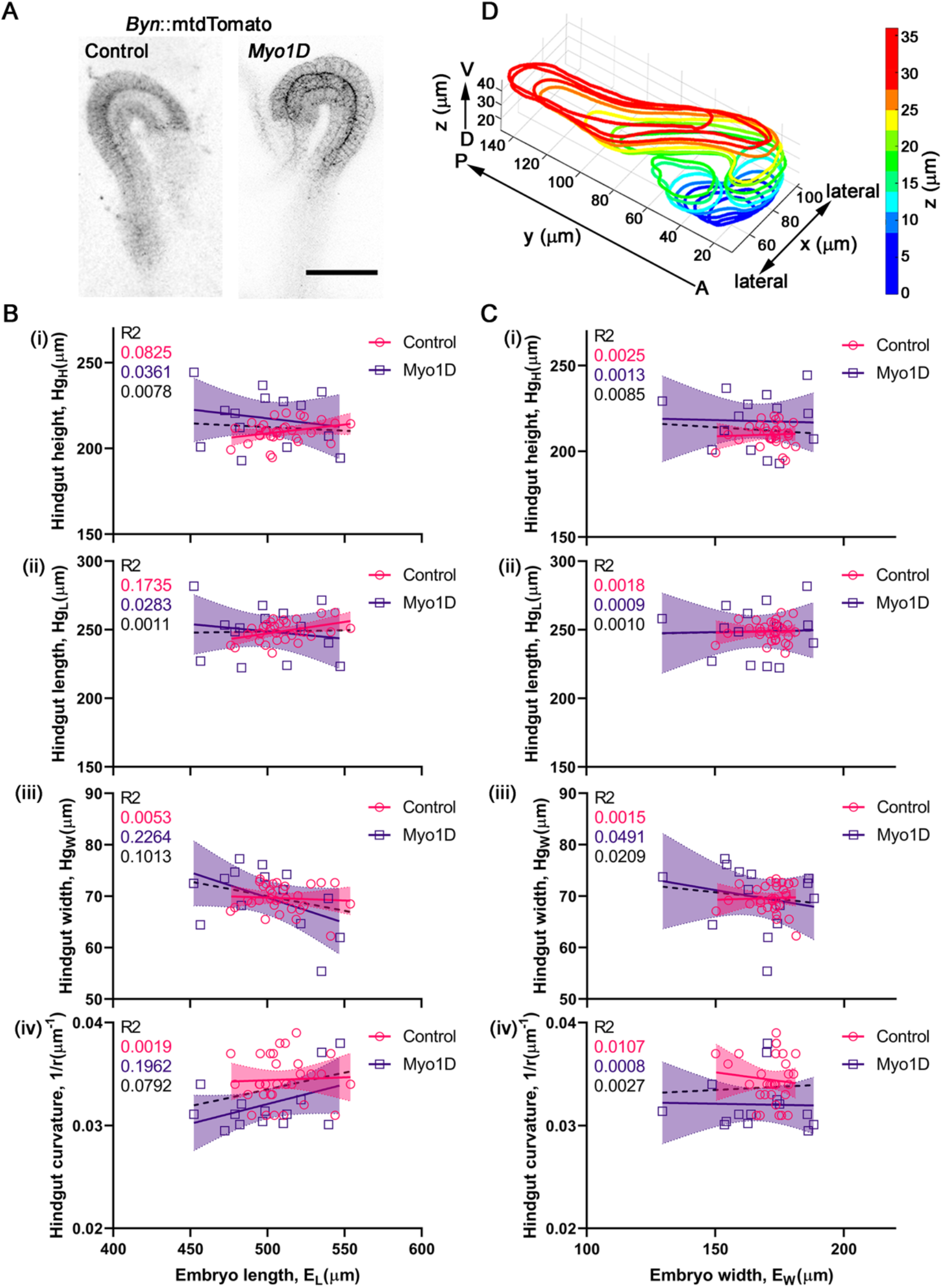
Hindgut scaling upon mechanical perturbation. A) Representative images of the hindgut in control and *Myo1D^-/-^* embryos, expressing *Byn*-GFP>UAS-myr::GFP. Note the inverted hindgut compared with control. B-C) Scaling of hindgut height Hg_H_ (i), total length Hg_L_ (ii), maximum width Hg_W_ (iii) and curvature (1/r) (iv) with embryo length (B) and width (C) in control (magenta) and mutant (purple) conditions. D) Three-dimensional segmentation of the hindgut in stage 14 (see also Movie S5). AP = Anterior-posterior axis, DV = Dorsal-ventral axis. Shaded regions represent 95% confidence interval on the linear fitting and r^2^ values given in the legend, colour coded by control (magenta), *Myo1D^-/-^* (purple) and all data (black). (n=31 for control and n=14 for *Myo1D^-/-^*), Scale bar = 50μm.

## Discussion

We have provided a dissection of scaling in three essential internal organs during *Drosophila* embryo development and shown that they display distinct scaling characteristics. The heart scales strongly with embryo length in stage 16 (Fig. 2). The hindgut shows weaker scaling but does display some adaptation to changes in embryo size, particularly in *TjGal4>fat2RNAi* embryos (Fig. 5). Lastly, the VNC length only weakly adjusts in size between embryos of significantly different morphology (Fig. 4). These results stand in stark contrast to the precision of gene networks early in the embryo [49, 50] where gene expression boundaries are closely scaled to embryo length. Our findings suggest that these gene boundaries are not translated into precisely scaled embryonic organs in general.

Our main observation is that the robustness of organ adjustment to changes in embryo size are quite distinct. Robustness in development is a longstanding problem given that many of the underlying processes are plastic in nature [51]. For example, under starvation, growth of the ovaries is delayed in the *Drosophila* larvae [52]. Conversely, the brain continues to develop similarly to healthy conditions for a substantially longer period [53–55]. The VNC is constructed of a highly stereotypic repetition of specific neurons and connections [34, 56]. Larvae from *TjGal4>fat2RNAi* embryos quickly reach similar absolute size and morphology to larvae from control embryos. This is likely due to the total volume of the hatching larvae not being substantially different, as the volume of the egg in *TjGal4>fat2RNAi* embryos is only decreased slightly compared to control embryos [27]. Therefore, the VNC may be sensitive to embryo volume, rather than specific size changes in one axis. This is plausible given the large size of the VNC prior to condensation. Overall, our results suggest that organs in the embryo adapt to specific changes in embryo size through organ-specific mechanisms.

Unlike most *Drosophila* organs, the embryonic heart is largely maintained throughout the fly life cycle [42]. Therefore, there may be advantages to ensuring the heart is correctly sized from an early stage of development as the organ is not substantially remodelled during either larval or pupal stages, limiting mechanisms that can adjust its size. We found that in heart scaling, the heart maintains cell number but changes cell size in smaller embryos. A range of organs regulate their size through modulating cell proliferation [5] or death [57, 58]. Instead, the heart cells appear to be mechanically malleable, enabling adjustment of size to fit within the constraining embryo environment. The mechanical structure of the heart cells has been analysed by electron microscopy [59], though which mechanical processes are driving the dynamic morphological changes during heart formation remains unknown.

The hindgut showed an intermediate response to changes in embryo size. In the hindgut, we see that the control embryos showed few characteristics of scaling but in the (wider) *TjGal4>fat2RNAi* embryos there is clear scaling behaviour (Fig. 5C-D). The surrounding organs may play an important role in restricting hindgut morphology. The midgut is positioned more anteriorly and may act to push against the hindgut as it extends. Further, the VNC lies beneath the hindgut, providing a barrier towards the ventral surface. Effectively, the surrounding organs may be acting to limit the available space for the hindgut and such limitations may also explain the observed shape changes in hindgut morphology between control and *TjGal4>fat2RNAi* embryos. In particular, the wider *TjGal4>fat2RNAi* embryos may facilitate the hindgut to extend further in its width. In adult flies, the gut size varies between males and females [60]. In future work, using our quantitative approach, we can explore whether these differences are apparent during embryogenesis. It will also be interesting to perturb the size of a specific organ and then explore how other organs adjust in size. Finally, we note that our analysis thus far has focused on the 2D projected hindgut morphology in the AP-lateral plane. However, the gut also extends in the dorsal-ventral (DV) axis (Fig. 6D and Movie S5). It is possible that the hindgut extends to differing degrees in the DV axis depending on constraints – for example, from neighbouring organs. However, it is currently challenging to dissect the 3D gut morphology at cellular resolution in time lapse movies and so we do not consider this further here.

It is interesting to compare these results with other organisms. Dorsal-ventral scaling in the early *Xenopus* embryo is regulated by embryo-size-dependent degradation [61]. By comparing different *Xenopus* species, evidence has also been found for transcriptional regulation of size [62]. The formation of the skull depends on neural crest cell migration. By varying where and when neural crest cells migrate to specific regions of the developing skull, the resulting size of the skull can vary substantially between avian species [63]. In mouse development, the formation of the pro-amniotic cavity depends on the size of the embryo [64]. In humans, there have been extensive studies of size regulation of organ growth during childhood [65]. The heart scales precisely with body mass during childhood [65]. In contrast, the brain does not scale with body size, with rapid early growth followed by much slower change in mass. In humans, how the embryo regulates organ size during embryonic development remains largely unknown. There appear to be a large range of mechanisms to ensure precise scaling in different organisms. Our results further suggest that even within the developing embryo the extent of organ scaling is potentially quite varied. Our study has focused on the embryonic stage of fly development, partly due to imaging accessibility for live, cellular-resolution imaging. Internal organ scaling may be differently regulated during larval and pupal stages, particularly given the large-scale remodelling processes that occur for the hindgut and VNC. Indeed, an interesting hypothesis is that the larval remodelling of organs may be able to correct for scaling errors from organ formation in the embryo.

Most previous work that has quantified scaling at a cellular level has focused on external organs, such as the wing. Organs such as the wing also grow to their final size. Here, we provide a detailed analysis of how three internal organs reach specific morphologies within the embryo given different embryonic constraints. These three organs have distinct mechanical behaviours resulting in different responses to changes in the embryo size. Understanding how organs scale and adapt to size changes remains a major challenge in development and this work suggests that we need new models to explain how tissues acquire specific morphologies when not simply growing to a final size.

## Acknowledgements

We thank Clarissa Halim, Mundzirah Bte Djuanda and Sham Tlili for data collection and preliminary data analysis. We thank Christopher Amourda, Christen Mirth, Nicholas Tolwinski and Yusuke Toyama for comments on the manuscript and members of the Saunders lab for constructive comments on the project. This work was funded by Mechanobiology Institute Seed Funding, the EMBO Global Investigator Program, and a Singapore Ministry of Education Tier 2 grant to T.E.S. (MOE2018-T2-2-138).

## Author Contributions

P.T. and T.E.S. designed the study. P.T. performed all experiments and analysis with assistance from H.R.. T.E.S. supervised the project and provided advice on statistical and image analysis. P.T. and T.E.S. analysed the results. P.T. and T.E.S. wrote the paper.

## Movies Legends

**Movie S1** Stereoscope movie showing *Drosophila* embryonic development from head involution until hatching in control and *TjGal4>fat2RNAi* embryos. There is no obvious difference in the morphological development except the embryo size. Related to Figure 1.

**Movie S2** Confocal time-lapse imaging showing migration and matching of the cardioblasts in control and *TjGal4>fat2RNAi* embryos. Cardioblasts marked by Hand::GFP. Apart from the cardioblasts, pericardial cells and visceral cells are also marked by Hand::GFP. Related to Figure 2.

**Movie S3** Confocal time-lapse imaging showing VNC condensation. VNC marked by ElavGal4>UAS-GFP in control and *TjGal4>fat2RNAi* embryos. Related to Figure 4.

**Movie S4** Confocal time-lapse imaging showing hindgut formation during development in control and *TjGal4>fat2RNAi* embryos. Hindgut marked by BynGal4>UAS-myr::GFP. Related to Figure 5.

**Movie S5** Z-stack of hindgut in a fixed stage 14 embryo labelled with BynGal4>UAS-myr::GFP (in green) and DE-Cad2 in red. This z-stack animation emphasises the 3D curved structure of the hindgut and in particular labels the hindgut lumen. Related to Figure 6D.

